# Evolution of a novel and adaptive floral scent in wild tobacco

**DOI:** 10.1101/738112

**Authors:** Han Guo, Nathalie D. Lackus, Tobias G. Köllner, Ran Li, Julia Bing, Yangzi Wang, Ian T. Baldwin, Shuqing Xu

## Abstract

Many plants emit diverse floral scents that mediate plant-environment interactions and attain reproductive success. However, how plants evolve novel adaptive floral volatiles remains unclear. Here, we show that in the wild tobacco, *Nicotiana attenuata*, a dominant species-specific floral volatile (benzyl acetone, BA) that attracts pollinators and deters florivore is synthesized by phenylalanine ammonia-lyase 4 (*NaPAL4*), isoflavone reductase 3 (*NaIFR3*), and chalcone synthase 3 (*NaCHAL3*). Transient expression of *NaFIR3* alone in *N. attenuata* leaves is sufficient and necessary for ectopic foliar BA emissions, and the BA emission level is increased by co-expressing *NaIFR3* with *NaPAL4* and *NaCHAL3*. Independent changes in transcription in all three genes contributed to intraspecific variations of floral BA emission. However, among species, the gain-of-expression in *NaIFR3* resulted in the biosynthesis of BA that was only found in *N. attenuata*. This study suggests that novel metabolic pathways associated with adaptation can arise via re-configurations of gene expression.

## Introduction

One of the major challenges in evolutionary biology is to understand the genetic mechanisms underlying the origin of phenotypic novelties. In flowering plants, floral volatiles are highly diverse and important for mediating ecological interactions between flowers and their visitors, including pollinators, florivores and pathogens^1,2^. In contrast to ubiquitous floral volatiles that are involved in the full spectrum of plant-pollinator interactions among different plant species^3^, species-specific floral volatiles likely evolved as the consequence of local adaption^4,5^. Although many of these species-specific floral volatiles are considered as novel adaptive traits in plants, how did they evolve remains largely unclear.

Benzyl acetone (4-phenylbutan-2-one; BA), a dominant nocturnal floral volatile in the wild tobacco species *Nicotiana attenuata*^6,7^, is known to attract hawkmoth pollinators such as *Manduca sexta* for outcrossing^8–10^ and simultaneously deter feedings from the florivore *Diabrotica undecimpunctata*^11^. Intriguingly, BA was not found in other *Nicotiana* species^12^, suggesting that BA is a species-specific floral volatile that underwent rapid evolution. Previous studies have shown that a chalcone synthase (*NaCHAL3*) is involved in the BA biosynthesis, as silencing this *NaCHAL3* resulted in reduced BA emissions in *N. attenuata*^8–10^. However, the biosynthetic machinery of BA and its evolution remain a mystery.

To identify genes involved in floral BA biosynthesis, we conducted QTL mapping, gene co-expression network analysis and genetic manipulations. We demonstrated that floral BA is synthesized from L-phenylalanine via three enzymes: phenylalanine ammonia-lyase 4 (NaPAL4), isoflavone reductase 3 (NaIFR3), and NaCHAL3. Comparative and evolutionary analyses among closely related species further suggest that the species-specific floral BA emission is resulted from a recent gain-of-expression in corolla limb in *NaIFR3*, a gene that originated before the divergence of *Nicotiana*. This study provides an example that novel metabolic pathways can arise via re-configuration the expression of existing genes.

## Results and discussion

### *NaPAL4* is required for the biosynthesis of BA in *N. attenuata* flowers

To identify the genetic basis underlying the variation in floral BA emissions, we performed quantitative trait loci (QTL) mapping using a *N. attenuata* advanced intercross recombinant inbred line (AI-RIL) population^13^, which was developed by crossing two inbred lines (AZ and UT) that differ in floral BA emissions (p = 0.0047, Figure 1A). By measuring the floral BA emission among individuals in the AI-RIL population, we identified one QTL locus on linkage group 5 (Figure 1B) that is significantly associated with floral BA emission. Because the genome of *N. attenuata* remains fragmented, to identify the candidate genes, we compared the corresponding genomic information of the identified QTL in *N. attenuata* and their syntenic genome annotations in *Petunia*. We found two homologous of *phenylalanine ammonia-lyase* (*PAL*) genes located at the corresponding QTL region. The PAL enzyme is involved in converting L-phenylalanine to *trans*-cinnamic acid (*t*-CA), which is the first step of most benzenoid metabolisms in *Petunia*^14^.

**Figure 1.**
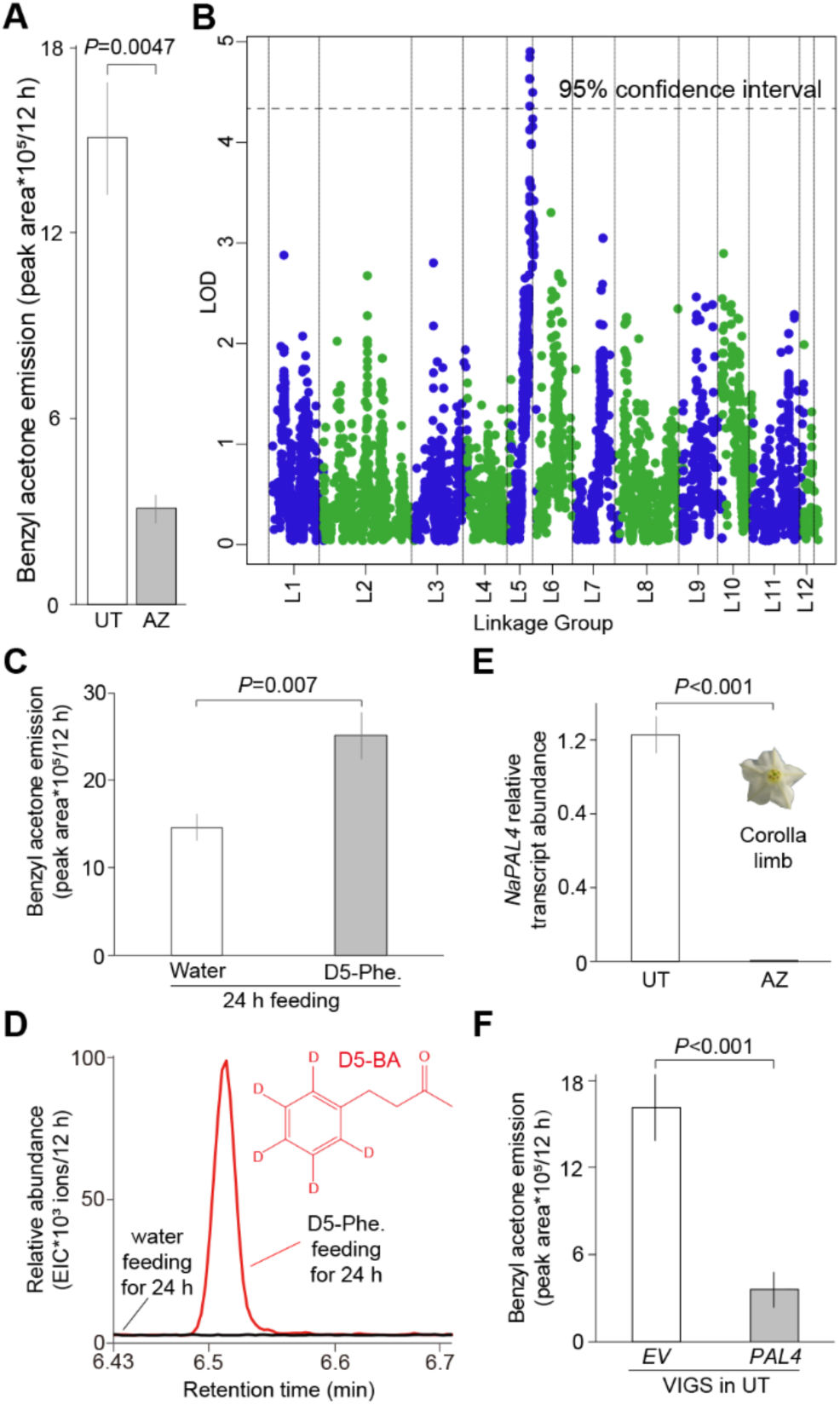
Phenylalanine ammonia-lyase 4 (*NaPAL4*) is involved in BA biosynthesis. (A) Quantitative differences in floral BA emission (mean ± SE, n = 8) between UT and AZ genotypes. (B) Floral BA emission is mapped to one QTL locus. The QTL locus on linkage group 5 is marked. The 95% confidence interval is indicated with a dashed line. LOD, log of the odds. (C) Significant differences of both nocturnal floral BA emission (mean ± SE, n = 8) (top panel) and deuterium_5_-BA (D5-BA) (bottom panel) are shown between water and L-Phenyl-d_5_-alanine (D5-Phe.) feeding on UT for 24 hours. Extracted-ion chromatogram, EIC. (D) Transcript abundance of *NaPAL4* (mean ± SE, n = 4) in corolla limb is different between UT and AZ genotypes. Corolla limb samples were harvested at 20:00. Transcript abundance was analyzed by RT-qPCR and relative to *elongation factor* gene in *N. attenuata* (*NaEF*). (E) In comparison to VIGS-*EV* plants, the levels of nocturnal floral BA emission (mean ± SE, n = 8) of VIGS-*NaPAL4* plants were significantly lower. For (A), (C) and (E), the trapping of floral BA was performed for 12 hours from 20:00 to 8:00. For (A), (C), (D) and (E), *P* values were calculated using Student’s-*t* tests.

To further examine whether BA is synthesized from L-phenylalanine, we stem-fed *N. attenuata* inflorescences (UT genotype) with deuterium-labeled phenylalanine (L-phenyl-d_5_-alanine) and measured floral BA emissions. In comparison to controls (water), the plants fed by L-phenyl-d_5_-alanine emitted significantly more BA (p = 0.007, Figure 1C). Furthermore, mass-spectrum analysis revealed that inflorescences feeding with L-phenyl-d_5_-alanine resulted in the occurrence of deuterium-labeled BA (Figure 1D), suggesting BA is synthesized from L-phenylalanine.

In the *N. attenuata* genome, we found four *NaPAL* candidates (*NaPAL1*-*4*). To examine the enzymatic activities of these NaPALs *in vitro*, we heterologously expressed *NaPAL1-4* in *Escherichia coli*. The results showed that all four NaPALs converted the substrate L-phenylalanine into *t*-CA *in vitro* (Figures S1A-S1F). We then compared the transcript abundance of these four candidates in the corolla limb, the tissue that is responsible for the emission of BA^6^, between the two parental lines that were used to generate the AI-RIL (UT and AZ). While the transcript abundance of *NaPAL1/2/3* in the corolla limb is similar between AZ and UT (Figures S1G-S1I), *NaPAL4* was only transcribed in UT but not in AZ (p < 0.001, Figure 1E). Further southern blot analysis showed that *NaPAL4* was absent in the AZ genome (Figures S1J and S1K).

To determine the function of *NaPAL4 in vivo*, we specifically silenced the expression of each *NaPAL* in *N. attenuata* UT plants using virus-induced gene silencing (VIGS) (Figure S1L) and measured their floral BA emission, respectively. Although *NaPAL1/2/3/4* could all convert L-phenylalanine into *t*-CA *in vitro*, only *NaPAL4*-silenced plants showed reduced floral BA emissions (p < 0.001, Figures 1F and S1M). Moreover, kinetic gene expression analysis of *NaPAL1-4* in eight different organs showed that *NaPAL4* had the highest transcript abundance in the corolla limb at ~ 20:00 (Figures S1N-S1Q), which is consistent to its role in BA biosynthesis. Additional analysis on the subcellular localization of NaPAL4 showed that NaPAL4 is localized to the endoplasmic reticulum membrane (Figures S1R-S1U), which is similar to many other phenylpropanoid biosynthesis-related PALs^15^. Taken together, these results suggest that the BA biosynthesis in *N. attenuata* requires *NaPAL4*-mediated conversion of L-phenylalanine to *t*-CA.

### *NaIFR3* is co-transcribed with *NaPAL4*, *NaCHAL3* and necessary for BA biosynthesis

Because *t*-CA has an extra carbon-carbon double bond in comparison to BA, we hypothesized that a reductase that removes the double bond is involved in the BA biosynthesis in *N. attenuata* (Figure 2A). To test this hypothesis, we first searched for the genes that are co-expressed with *NaPAL4* using our previously established *Nicotiana attenuata* datahub platform^16^. In addition, since previous study showed that a chalcone synthase, *NaCHAL3* (renamed from *NaCHAL1* to *NaCHAL3* according to phylogenetic analysis in this study), is involved in the BA biosynthesis^8^, we also included *NaCHAL3* in the gene co-expression analysis. Among all co-expressed genes, *NaIFR3*, which structurally belongs to the family of NADPH-dependent reductases^17^, showed a similar corolla limb-specific expression to both *NaPAL4* and *NaCHAL3* (Figure 2B). A kinetic of the transcript abundance showed that *NaPAL4*, *NaIFR3* and *NaCHAL3* are all highly transcribed in the night (Figure 2C), which is consistent to the nocturnal floral emission of BA^6,11^.

**Figure 2.**
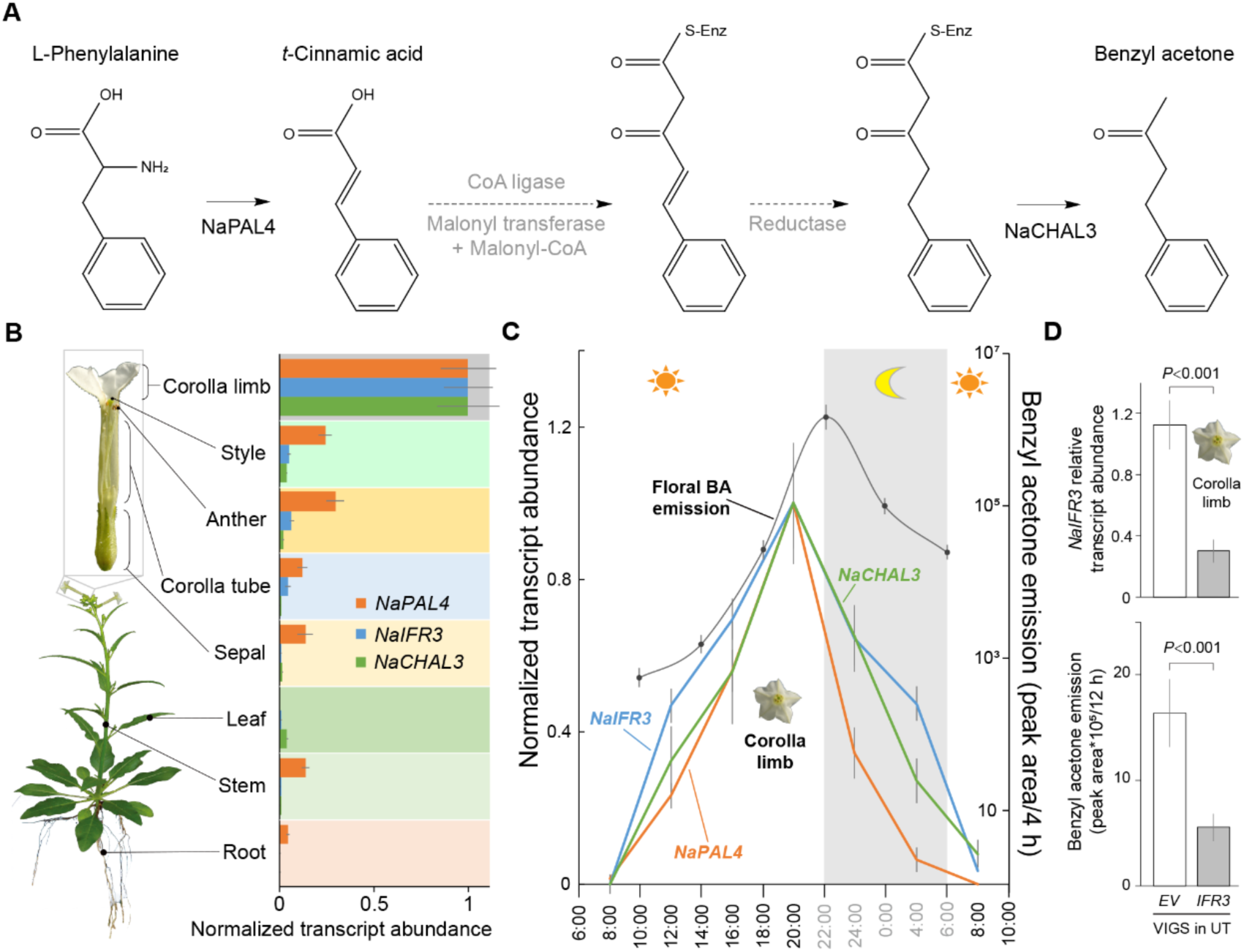
*NaIFR3* is co-transcribed with *NaPAL4* and *NaCHAL3* and involved in BA biosynthesis. (A) Predicted biosynthesis pathway of BA. Dashed lines with arrow heads indicate the putative steps and the putative corresponding enzymes are shown in grey. (B) *NaPAL4*, *NaIFR3* and *NaCHAL3* (mean ± SE, n = 4) are co-transcribed abundantly in corolla limb. Transcript abundance was analyzed by RT-qPCR. Different tissues were harvested at 20:00. The transcript abundance of each gene is relative to *NaEF* and normalized (normalized as X’ = X/X_max_) among different tissues. (C) Emission kinetics of floral BA (mean ± SE, n = 8) are consistent to kinetics of *NaPAL4*, *NaIFR3* and *NaCHAL3* transcription in corolla limb (mean ± SE, n = 4). The trapping of floral BA was performed for the periods of 4 hours, e.g. 8:00-12:00, 12:00-16:00 and so on. Corolla limb samples were harvested every 4 hours, e.g. 8:00, 12:00 and so on. Transcript abundance of each gene was analyzed by RT-qPCR and relative to *NaEF* and normalized (normalized as X’ = X/X_max_) among different time points. (D) In comparison to VIGS-*EV* plants, the levels of nocturnal floral BA emission (mean ± SE, n = 8) of VIGS-*NaIFR3* plants were significantly lower. The trapping of floral BA was performed for 12 hours from 20:00 to 8:00. *P* values were calculated using Student’s-*t* tests.

To examine the function of *NaIFR3 in vivo*, we specifically silenced the expression of *NaIFR3* in *N. attenuata* (UT) using VIGS (p < 0.001, Figures 2D top panel and S2A) and measured the floral BA emission. The silencing of *NaIFR3* did not result in any morphological changes of the flowers, but specifically reduced the floral BA emission in *N. attenuata* (p < 0.001, Figure 2D bottom panel).

We then compared the transcript abundance of *NaIFR3* between AZ and UT in corolla limbs. Interestingly, while *NaIFR3* was highly transcribed in UT, it is only transcribed at the basal level in AZ (p < 0.001, Figure S2B). Further southern blot analysis showed that the low transcript abundance of *NaIFR3* in AZ is not because of a gene loss (Figure S2C). Because only one locus was found in the QTL mapping, it seems likely that *NaPAL4* and the *cis-*or *trans*-regulator that resulted in differences in the transcript abundance of *NaIFR3* are co-located.

Due to the difficulties to obtain a stable substrate, it is challenging to directly examine the biochemical function of *NaIFR3 in vitro*. Therefore, we directly examined the function of *NaIFR3* by ectopically expressing *NaIFR3*, *NaPAL4* and *NaCHAL3* in *N. attenuata* leaves, either individually or in combinations. At 48 hours after transformation, ectopic transcript abundance of *NaPAL4*, *NaIFR3* and *NaCHAL3* and foliar BA emission were measured in the transformed *N. attenuata* rosette leaves. The results showed that ectopic expression of *NaIFR3* alone is already sufficient for low-levels of foliar BA emission (Figure 3A), suggesting that other components of BA biosynthesis already exist in *N. attenuata* leaves. Consistently, in *N. attenuata* leaves, we found a relatively high transcript abundance of *NaPAL1* and *2* (Figures S1N and S1O) both showing *in vitro* the ability to convert L-phenylalanine into *t*-CA (Figures S1C and S1D) and we observed transcripts of *NaCHAL3* (Figures 2B and S2K), albeit in low abundance. Further co-expression of *NaIFR3* with *NaPAL4* and *NaCHAL3* in *N. attenuata* leaves, either in pairwise combinations or all three together, significantly increased BA emission in comparison to expressing *NaIFR3* alone (Figure 3A). These results revealed that *NaIFR3* is required for BA biosynthesis in *N. attenuata* and co-expression of *NaPAL4*, *NaIFR3* and *NaCHAL3* in leaves is sufficient for ectopic foliar BA emission.

**Figure 3.**
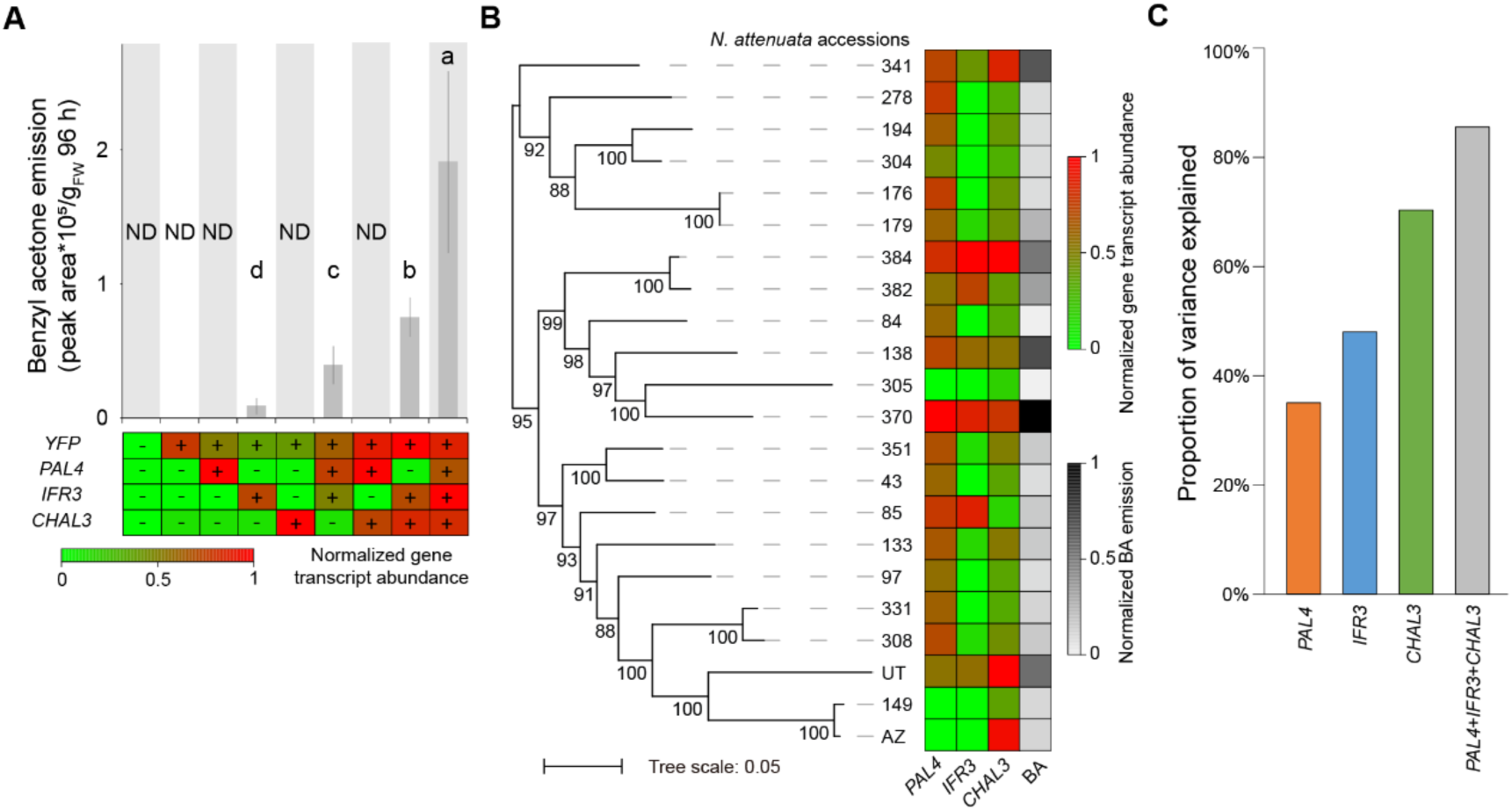
Ectopic expression of *NaIFR3* is sufficient for foliar BA emission and transcriptional changes of *NaPAL4*, *NaIFR3* and *NaCHAL3* contributed to the intraspecific variations of floral BA emission. (A) Foliar BA emission (mean ± SE, n = 8) (top panel) with corresponding ectopic transcription of *NaPAL4*, *NaIFR3* and *NaCHAL3* (bottom panel) are shown. The trapping of foliar BA was performed for 96 hours after transformation. Different letters indicate significant differences in a Tukey-corrected *post-hoc* test following a one-way ANOVA (*P* < 0.05). ND: Not detected. The complete open reading frames of *NaPAL4*, *NaIFR3* and *NaCHAL3* were fused downstream of *YFP* and transiently expressed in *N. attenuata* rosette leaves, individually or in different combinations together. Heatmap (bottom panel) representing the normalized transcript abundances of *NaPAL4*, *NaIFR3* and *NaCHAL3* (relative to *NaEF* and normalized as X’ = X/X_max_ among different transformations), respectively. + or -, presence or absence of the respective transgene. (B) Heatmap representing the transcript abundances of *NaPAL4*, *NaIFR3*, *NaCHAL3* and nocturnal floral BA emission in 22 *N. attenuata* natural accessions. The phylogenetic tree of the 22 accessions on left panel was built based on their genomic sequences. Numbers on branches indicate the bootstrap percentage values calculated from 100 replicates, and only values greater than 50% are shown. The transcript abundance is relative to *NaEF* and normalized as X’ = X/X_max_ among different accessions. The trapping of floral BA was performed for the 12 hours from 20:00 to 8:00. (C) Relationship between floral BA emission and transcript abundance of *NaPAL4*, *NaIFR3* and *NaCHAL3* in 22 natural accessions. The y-axis denotes the proportion of BA variation that could be explained by the changes of genes’ transcription.

### Independent expression changes of *NaPAL4*, *NaIFR3* and *NaCHAL3* resulted in intraspecific variations of floral BA emission

To further investigate the genetic mechanisms underlying natural variations of floral BA emission in *N. attenuata*, we measured the transcript abundance of *NaPAL4*, *NaIFR3* and *NaCHAL3* and the floral BA emission among 22 natural accessions (Table S3) that had been re-sequenced with low coverage. Overall, the variations in floral BA emission and the transcript abundance of *NaPAL4, NaIFR3 and NaCHAL3* did not show a clear correlation with their genetic distance that was calculated using genome-wide SNPs (Figure 3B). This suggests that the variations of floral BA emission were not a result from historic demographic changes of *N. attenuata*, but likely due to variations of local adaptations to pollinators or florivores^18^. Furthermore, expression changes in *NaPAL4, NaIFR3 and NaCHAL3* were also not correlated among genotypes, indicating that changes in the expression among these genes were largely independent.

We then estimated the extent to which the expression changes of each of the three genes contributed to the natural variation of floral BA emissions in *N. attenuata*. The results showed that the changes in the transcript abundance of *NaPAL4*, *NaIFR3*, *NaCHAL3* and all three genes together could explain ~38%, ~50%, ~70% and ~85% of the floral BA emission variance among the 22 accessions, respectively (Figure 3C). In contrast, variations in *NaPAL1*, *2*, *3* and *1-3* together, which are likely not involved directly in the floral BA biosynthesis, can only explain ~11%, ~5%, ~10% and ~15% of the floral BA emissions, respectively (Figure S2E). Together, these results suggest that the intraspecific variations in the floral BA emission in *N. attenuata* largely resulted from independent changes in the expression of each of its biosynthetic genes.

### Gain-of-expression in *NaIFR3* resulted in the BA biosynthesis in *N. attenuata*

Based on both the *in vivo* functions and the putative enzymatic activities of *NaPAL4*, *NaIFR3* and *NaCHAL3*, we derived a possible BA biosynthesis pathway in *N. attenuata* (Figure S3A). It is also possible that *NaCHAL3* might act earlier than *NaIFR3* in the pathway. However, a previous study in *Rheum palmatum* showing that the benzalacetone synthase (BAS), which shares 70% amino acid sequence similarity to *NaCHAL3*, catalyzes the one-step decarboxylative condensation of 4-coumaroyl-CoA with malonyl-CoA to produce a diketide benzalacetone^19,20^. Therefore, it is likely that NaCHAL3 is the enzyme that is responsible for the final product of BA emission.

To investigate the evolution of floral BA biosynthesis in *N. attenuata*, we first performed phylogenomic analysis using the available genomic data and analyzed the evolutionary history of *NaPAL4*, *NaIFR3* and *NaCHAL3*. The results showed that *NaPAL4* originated before the whole genome triplication (WGT) that was shared among Solanaceae species (Figures 4A and S3B), while *NaIFR3* and *NaCHAL3* originated from gene duplications that are specific to the *Nicotiana* genus (Figures 4A, S3C and S3D). The duplicated copies of *NaPAL*, *NaIFR* and *NaCHAL* all showed expression divergence, both among tissues and in their temporal dynamics (Figures S1N-S1Q and S2F-S2K). Together, these results suggest that the BA biosynthesis resulted from recruiting both ancient and recent duplicated genes.

**Figure 4.**
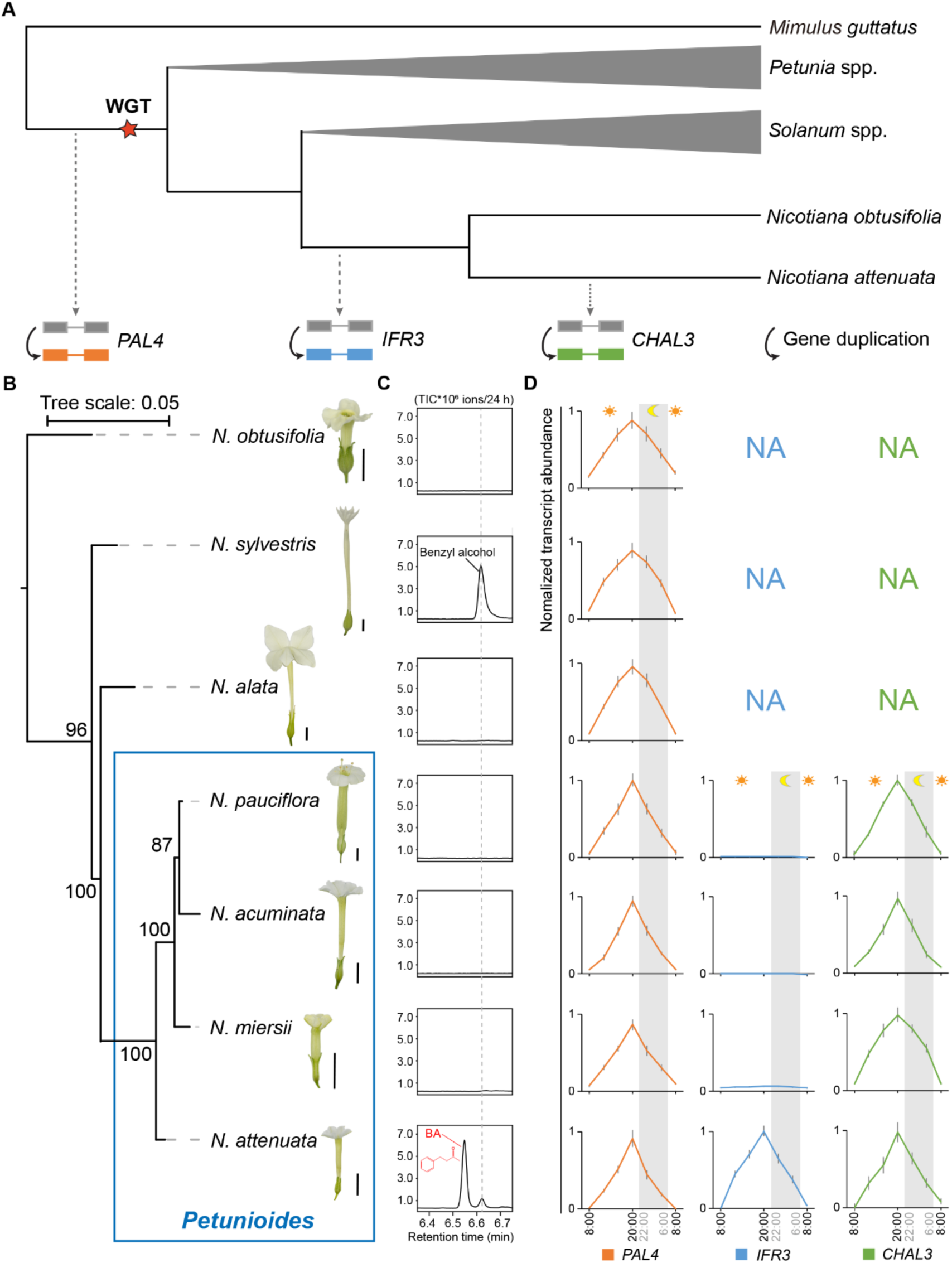
Gain of corolla limb expression of *NaIFR3* coincides with floral BA biosynthesis in *N. attenuata*. (A) Schematic evolutionary history of *NaPAL4*, *NaIFR3* and *NaCHAL3*. WGT, whole genome triplication. Dashed lines with arrow heads indicate the estimated time points of corresponding gene duplication. (B) The phylogenetic tree of the seven *Nicotiana* species was built based on partial nepGS sequences. Numbers on branches indicate the bootstrap percentage values calculated from 1000 replicates, and only values greater than 50% are shown. Scale bars of flowers: 1 cm. (C) Floral emission of BA is species-specific in *N. attenuata*. The trapping of floral volatile was started at 8:00 and was performed for 24 hours. Total ion current, TIC. (D) Kinetic of the transcript abundance of *NaPAL4* (left panel), *NaIFR3* (middle panel) and *NaCHAL3* (right panel) in corolla limbs among seven *Nicotiana* species. Transcript abundance (mean ± SE, n = 4) was analyzed by conserved primers and RT-qPCR. The transcript abundance of each gene is relative to *NaEF* and normalized (normalized as X’ = X/X_max_) among different species in kinetics. When the gene could not be identified from either genomic data or homologous cloning, we labeled the expression as NA (not available).

Because *N. sylvestris* is the most closely related species to *N. attenuata* that has its genomic sequence available, phylogenomic analysis is limited to the time before speciation between *N. sylvestris* and *N. attenuata* (Figure 4A). To gain more insights into the evolution of BA biosynthesis after this speciation event, we further analyzed the floral volatiles in seven closely related *Nicotiana* species (Figures 4B and 4C). Consistent with a previous study^12^, BA was only found in *N. attenuata* (Figure 4C).

We amplified and sequenced the cDNA of *IFR* and *CHAL* among the seven *Nicotiana* species. Phylogenetic analyzes of the cDNA sequences of *IFR* and *CHAL* among these closely related species were consistent to the analysis based on the genomic sequences: *NaIFR3* originated in the ancestor of *N. attenuata* and *N. obtusifolia* (~12.5MYA)^21^ and *NaCHAL3* occurs specifically in the clade of *Petunioides* (~9.1 MYA)^22^.

We further analyzed the expression kinetics of *NaPAL4*, *NaIFR3* and *NaCHAL3* in the corolla limb of all seven species. Interestingly, like in *N. attenuata*, *NaPAL4* showed a high transcript abundance at 20:00 among all species (Figure 4D) and *NaCHAL3* was highly transcribed at 20:00 among the four species of *Petunioides*. This indicates that *NaPAL4* and *NaCHAL3* might be involved in other floral metabolic pathways in the different *Nicotiana* species. However, although *NaIFR3* existed in all four species of *Petunioides*, it was only transcribed in *N. attenuata* corolla limbs and showed evening-specific expression pattern. Because *NaIFR1/2*, the ancestor/homologs of *NaIFR3*, were expressed either in styles, stems or leaves, the corolla limb expression of *NaIFR3* is likely due to a tissue-specific gain-of-expression event. The event either specifically occurred in *N. attenuata* or in the ancestor of *Petunioides* then only maintained in *N. attenuata*. Together, these results suggest that tissue-specific gain-of-expression in an existing gene, *NaIFR3*, have resulted in the evolution of the BA biosynthesis that mediates both pollinator attraction and florivore deterrence.

In summary, this study demonstrated that a new adaptive metabolic pathway in plants can arise from expression changes in a single gene. Such mechanism underlying the emergence of new metabolic pathways that mediate key ecological interactions might not only explain the evolution of amazing diversity of specialized metabolites in plants^23,24^, but also highlight the potential for breeding eco-friendly crops via metabolic engineering.

## Methods

### Plant material

The *N. attenuata* Utah (UT) and Arizona (AZ) wild type seeds were originally collected from plants growing in a large natural population near Santa Clara, Utah, USA^25^, and a 20-plant population near Flagstaff, Arizona, USA^26^. They were inbred for 31 and 22 generations, respectively, in the glasshouse. Seeds of the G2 accession were collected in Utah as described in^27^. Additional natural accessions were collected by Ian T. Baldwin throughout the southwestern United States and inbred for one generation in the glasshouse^28^. To develop the AI-RIL population, UT and AZ were first crossed to generate F1 plants, which were then self-fertilized to generate 150 F2 plants. From F2 to F6, in each generation, we intercrossed ~150 progeny using a random mating and equal contribution crossing design^29^. For generation F7, two seeds from each of the crosses at F6 were germinated and used for the single-seed descendent inbreeding process. In total, five generations of inbreeding were conducted.

### Plant growth

All seeds were germinated following the protocol described by Krügel *et al.* (2002)^30^. Plants were grown under in glasshouse conditions (26 ± 1°C; 16h: 8h, light: dark)^30^. For the VIGS and transient transformation assay, plants were grown in a climate chamber under a constant temperature of 26 °C and 16h:8h (light: dark) light regime, and 65% relative humidity^31^.

### Sampling of floral and foliar BA emissions

We measured floral BA from flowers of *Nicotiana* plants (~50 days after germination). Flowers of *Nicotiana* species remain open for three days and the flower age affects the quantity of floral volatiles^32^. Therefore, we removed all open flowers in the morning (7:00-9:00) of the day of volatile trapping. For each biological replicate, one freshly opened flower was taken out of the plant at 20:00 and incubated with polydimethylsiloxane (PDMS) tubes in a sealed 8 mL glass vial (MACHEREY-NAGEL)^33^.

To measure foliar BA emission, a transformed rosette leaf of *N. attenuata* was taken and incubated with PDMS tubes in a sealed 8 mL glass vial (MACHEREY-NAGEL).

### Volatile analysis by TD-GC-MS

PDMS tubes were placed in an autosampler with thermal desorption unit (TDU; TD-20, Shimazu), which was connected to a quadruple GC-MS (QP-2010-Ultra, Shimazu) for analysis. Specifications for desorption conditions, the used columns and the spectra reading and identification, were as described by Kallenbach *et al.*^33^ and Schuman *et al.*^34^. For all of the volatiles, a 1:100 split was used to avoid overloading the detector.

### Isotope-labeled phenylalanine feeding assay

Shoots with mature flower buds from UT plants (50 days after germination) were cut, inserted into a falcon tube (50 mL) and fed with either 10 mM L-phenyl-D5-alanine (Sigma-Aldrich, Cat# 615870) or water for 24 hours, respectively.

### Heterologous expression and enzyme assays of NaPAL1-4

The *E. coli* strain BL21 Star (DE3) (Thermo-Fisher) was used for expression of the complete open reading frames of *NaPAL1-4.* Since expression of native *NaPAL4* yielded no protein, a codon optimized version was synthesized and used for enzyme characterization. Cultures were grown at 37°C, induced at an OD_600_ = 0.6 with 1 mM IPTG, subsequently placed at 18°C, and grown for another 20 hours. The cells were collected by centrifugation and disrupted by freezing in liquid nitrogen and following thawing (five times) in chilled extraction buffer (50 mM Tris-HCl, 500 mM NaCl, 20mM Imidazole, 10% Glycerol; 1% Tween20; pH8,6). Cell fragments were removed by centrifugation at 14,000 g, the supernatant was purified with HisPur Cobalt Resin (Thermo-Fisher), and the purified proteins were concentrated with Amicon Ultra-0.5 Centrifugal Filter devices (Merck Millipore) following manufactures instructions. To determine the catalytic activity of recombinant PAL enzymes, assays containing 20 µl of the purified protein, 79 µl assay buffer (50 mM Tris-HCl, 500 mM NaCl, 10% Glycerol; pH8,6) and 1 mM phenylalanine as substrate were incubated for 3 hours at 35°C. Reaction products were analyzed using LC-MS/MS.

### LC-MS/MS analysis of NaPAL enzyme products

Chromatography was performed on an Agilent 1260 Infinity II HPLC system (Agilent Technologies). Separation was achieved on a Zorbax Eclipse XDB-C18 column (50 × 4.6 mm, 1.8 µm, Agilent). As mobile phases A and B, formic acid (0.05%) in water and acetonitrile were employed, respectively, with a mobile flow rate of 1.1 mL/min. The elution profile was: 0-0.5 min, 10% B; 0.5-4.0 min, 10-90% B; 4.0-4.02 min 90-100% B; 4.02-5.50 min 100% B; 5.51-8.00 min 10% B. The column temperature was set at 20°C. The liquid chromatography was coupled to an API-6500 tandem mass spectrometer (Sciex) equipped with a Turbospray ion source (ion spray voltage, −4500 eV; turbo gas temperature, 650 °C; nebulizing gas 60 psi., heating gas 60 psi, curtain gas 45 psi, collision gas medium). Measurements were performed in negative mode. Multiple reaction monitoring (MRM) was used to monitor a parent ion → product ion reaction for the PAL substrate phenylalanine (*m/z* 164 → 147.0, CE −18 V, DP −50 V) and the reaction product cinnamic acid (*m/z* 147 → 103.0, CE −16 V, DP −50 V). Analyst 1.6.3 software was used for data processing and analysis (AB Sciex).

### Virus-induced gene silencing, VIGS

Leaves of young *N. attenuata* plants were agroinfiltrated with *pBINTRA* and *pTV-NaPAL1/2/3/4* or *pTV-NaIFR3* according to a published protocol optimized for VIGS in *N. attenuata*^35^. Plants co-infiltrated with *pBINTRA* and *pTV00* were used as control. All VIGS experiments were repeated at least three times.

### Southern blot analysis

A total amount of 20 µg genomic DNA was digested overnight at 37°C with 100 U *Eco*RV or *Xba*I or *Bam*HI or *Hin*dIII (New England Biolabs) in independent reactions. The digested DNA was separated on a 0.8% (w/v) agarose gel for 15 h at 30 Volt. DNA was blotted overnight onto a Gene Screen Plus Hybridization Transfer Membrane (Perkin-Elmer) using the capillary transfer method. For hybridizations, gene specific fragments that were used for VIGS (primer pairs listed in **Table S1**) were radiolabeled with [α-32P] dCTP (Perkin-Elmer) using the Rediprime II DNA Labeling System (GE Healthcare) according to the manufacturer’s instructions. The blot was washed twice at high stringency (0.1× SSC and 0.5% SDS for 20 min). Membranes hybridized with radioactive probes were exposed for 12 hours to a phosphor screen (FUJIFILM imaging plate, BAS-IP MS 2340) in FUJIFILM BAS cassette 2340. Then the phosphor screen was scanned by Fujifilm FLA-3000 fluorescence laser imaging scanner for visualization.

### Transient transformation for subcellular localization analysis and ectopic expression in leaves

The construction of 35S::*YFP*, 35S::*YFP-NaPAL4*, 35S::*YFP-NaIFR3* and 35S::*YFP-NaCHAL3* reporter fusions were carried out as described by Earley *et al*.^36^. The open reading frame encoding these genes were amplified and introduced into intermediate pENTR plasmid (Thermo-Fisher, Cat# K240020) and then introduced into pEarleyGate 104 to generate YFP fusion constructs. The used primers are listed in **Table S1**. Recombined plasmids were then transformed into *Agrobacterium tumefaciens* strain GV3101 for subsequent plant transformation. Leaves of 3-weeks old *N. attenuata* plants were co-infiltrated with *A. tumefaciens* cells containing different plasmids. To detect the localization of NaPAL4, 35S::*XA10-CFP* was co-transformed with *NaPAL4* and to generate endoplasmic reticulum (ER)-specific CFP fluorescent signal^37,38^. Fluorescence was visualized 48 h following the inoculation with a Zeiss LSM 510 Meta confocal microscope (Carl Zeiss).

### Phylogenetic tree construction

The phylogenetic relationship among 22 *N. attenuata* accessions was constructed using genome-wide SNP data. In brief, each accession was sequenced in low coverage (5-10 X) using Illumina HiSeq 2000 (pair-end). The short reads were mapped to *N. attenuata* reference genome^21^ using BWA-mem^39^. Genome-wide variants were called using GATK pipeline. VCFtools^40^ was used to remove non biallelic variants, reads coverage less than 1 or greater than 1000, missing data in more than 40% of accessions, minimum SNP quality less than 30, mapping quality less than 50 and indels. BCFtools^41^ was used to prune SNPs in linkage (if two SNPs in a 1000kb window have their r^2^ > 0.5 were discarded). This resulted in 157,833 high quality SNPs. These SNPs were used for building phylogenetic tree using RAxML-NG (v0.9.0)^42^ with 100 bootstraps. The best tree inference model “TVM+G4” was estimated by modeltest-NG^43^. To construct the phylogenetic tree of *PAL4*, *IFR3* and *CHAL3*, we used the Fishing Gene Family pipeline ^44^ with minor modifications. In brief, the protein sequence of each of the three gene was used as the bait and genomic sequences of different Solanaceae species were used as the database. The extracted exon sequences were then aligned using GeneWise and the phylogenetic tree was constructed using PhyML (v3.3.3)^45^. The phylogenetic tree of seven *Nicotiana* species was constructed using PhyML(v3.3.3)^45^ based on partial nepGS gene sequences obtained from Clarkson *et al.* 2010^46^. Visualization of phylogenetic tree were conducted by iTOL v4.4.2^47,48^.

### QTL mapping

The genotype information of all AI-RIL plants and the linkage map were obtained from the dataset reported earlier^13^. The R package QTLRel was used for QTL mapping following the tutorial^49^. Briefly, the relationship among different individuals was first estimated based on pedigree information. The peak area of each compound was log-transformed. Samples with missing genotype or phenotype information were removed. In total, 207 samples were used for QTL mapping. Then the variance of the traits within the population was estimated via “estVC” and missing information of the genotypes was imputed using the function “genoImpute”. The estimated trait variance and imputed genotypes were then used for the genome-wide scan. The empirical threshold was estimated based on 500 permutations. The additive and dominant effects of the candidate QTLs were estimated by fitting a multiple QTL model using the function “gls”.

### Quantitative RT-PCR

Total RNA was isolated using the RNeasy Plant Mini Kit (QIAGEN, Cat#74903), and 1000 ng of total RNA were reverse transcribed using the PrimeScript RT-qPCR Kit (TaKaRa, Cat#RR037B). At least four independent biological replicates were collected and analyzed. RT-qPCR was performed on the Stratagene 500 MX3005P using a SYBR Green reaction mix (Eurogentec, Cat#10-SN2X-03T). The primers used for mRNA detection of target genes by RT-qPCR are listed in **Table S1**. The mRNA of *N. attenuata elongation factor* (*NaEF*) was used as internal control.

### Proportion of BA emission variance explained by gene expression levels among natural accessions

To analyze the correlation between floral BA emission and transcripts level of the three candidate genes among 22 *N. attenuata* natural accessions, we firstly applied square root transformation followed by Z-Score normalization with “scale” function in R (https://cran.r-project.org). Then linear regression models were used to fit the transformed data. We used floral BA emission as response variable and transcript abundances of each gene as independent variable using the “lm” function in R.

### Data availability

The data generated or analyzed during the current study are included in this published article (and its Supplementary Information) or are available from the corresponding author on reasonable request.

## Acknowledgments

We thank the glasshouse team of the Max Planck Institute for Chemical Ecology for plant cultivation; Dapeng Li, Wenwu Zhou, Klaus Gase and Eva Rothe for technical assistance; Lei Hou, Rayko Halitschke, Danny Kessler and Martin Schäfer for fruitful discussions. This work was supported by funding from the Swiss National Science Foundation (PEBZP3-142886 to S.X.), a European Commission Marie Curie Intra-European Fellowship (IEF) (328935 to S.X.), the Max Planck Society, and the Advanced Grant no. 293926 of the European Research Council to I.T.B.

## Author contributions

Conceptualization, S.X and H.G; QTL mapping and gene candidate validation, S.X. and H.G.; intra- and interspecific gene expression analysis, VIGS, isotope-labeled phenylalanine feeding and Southern blot assay, H.G.; transient transformation for subcellular localization analysis and ectopic expression in leaves, H.G. and R.L.; heterologous expression and enzyme assays of NaPAL1-4, H.G., N.D.L. and T.G.K.; sampling of floral and foliar BA emissions and volatile analysis by TD-GC-MS, H.G. and J.B.; phylogenetic analysis, S.X., H.G. and Y. W.; writing – original draft, S.X. and H.G.; funding acquisition, S.X. and I.T.B.; resources, S.X. and I.T.B.; supervision, S.X.

## Declaration of interests

The authors declare that there is no conflict of interest regarding the publication of this article.

## Additional information

Correspondence and requests for resources and reagents should be addressed to S.X. Requesting *N. attenuata* seeds should be addressed to I.T.B.

