## Supplementary information for "Evolution of a novel and adaptive floral scent in wild tobacco"

**Supplemental figures and legends**

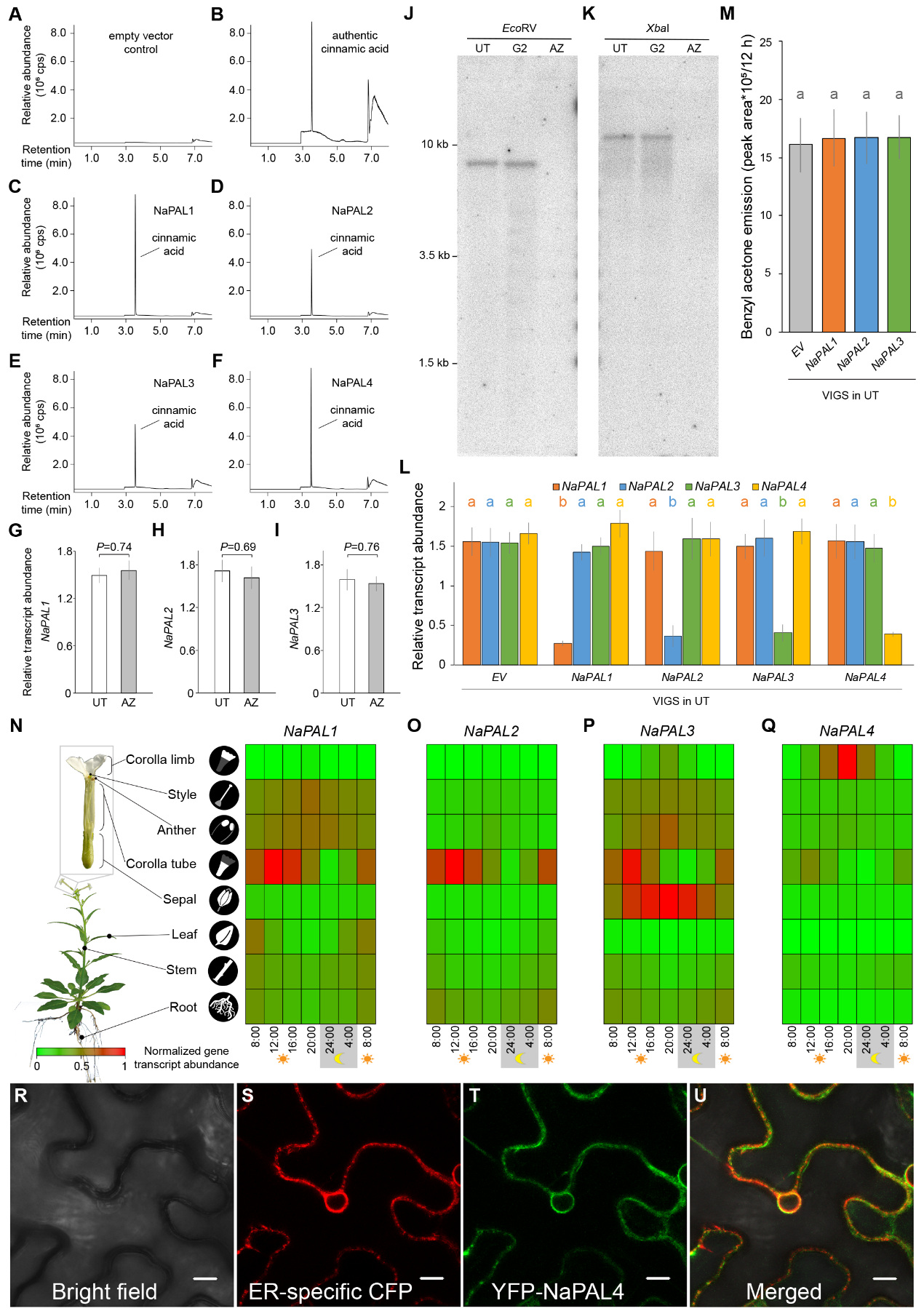

**Figure S1. Function and expression of NaPAL1-4.**

(A) to (F), Enzymatic functions of NaPAL1-4. The genes were heterologously expressed in *Escherichia coli* and the affinity-purified proteins were incubated with phenylalanine as substrate. Enzyme products were analyzed using liquid chromatography-tandem mass spectrometry. Cinnamic acid formation was verified using an authentic standard. An empty vector control showed no PAL activity. Counts per second, CPS.

(G) to (I), Transcript abundance of *NaPAL1* (G), *NaPAL2* (H) and *NaPAL3* (I) (mean ± SE, n = 4) in corolla limbs is not significantly different between UT and AZ genotypes. Corolla limb samples were harvested at 20:00. Transcript abundance was analyzed by RT-qPCR and relative to *elongation factor* gene in *N. attenuata* (*NaEF*). *P* values were calculated using Student’s-t tests.

(J) and (K), Genomic DNA (20 μg) from three phenotypes of plants (genotype G2 emitting abundant BA as another positive control) were digested with *Eco*RI (J) or *Xba*I (K). After separation by agarose gel electrophoresis and blotting, the DNA was hybridized with NaPAL4 partial cDNA (1596-1845 bp, 250 bp). The blot was washed at high stringency (0.1× SSC and 0.5% SDS for 20 min twice).

(L) Silencing the transcription of *NaPAL1*-*4* by VIGS in corolla limb. Corolla limb samples were harvested at 20:00. Transcript abundance (mean ± SE, n = 4) were analyzed by RT-qPCR and is shown relative to *NaEF*. Different letters indicate significant differences in a Tukey-corrected *post-hoc* test following a one-way ANOVA (*P* < 0.05).

(M) There is no significant difference in the nocturnal floral BA emission between VIGS-*EV* and VIGS-*NaPAL1*-*3* plants. The trapping of floral BA was performed for the 12 hours from 20:00 to 8:00. Same letter indicates no significant difference in a Tukey-corrected *post-hoc* test following a one-way ANOVA (*P* < 0.05).

(N) to (Q), Heatmap representing the kinetic of the transcript abundance of *NaPAL1* (N), *NaPAL2* (O), *NaPAL3* (P) and *NaPAL4* (Q) among eight tissues of the UT genotype. The transcript abundance is relative to *NaEF* and normalized as X’ = X/X_max_ among different tissues and different time points.

(R) to (U), The complete open reading frame of *NaPAL4* was fused downstream of *YFP* and transiently expressed in *N. attenuata* rosette leaves. (R) bright field image of the leaf area; (S) The location of the endoplasmic reticulum was determined by the fluorescence of the ER-specific fusion protein XA10-CFP (red); (T) YFP fluorescence indicates the location of the fusion protein YFP::NaPAL4 (green); (U) Merged signals of both the CFP and YFP fluorescence emitted from the leaf area. Scale bar in (R) to (U): 10 μm.

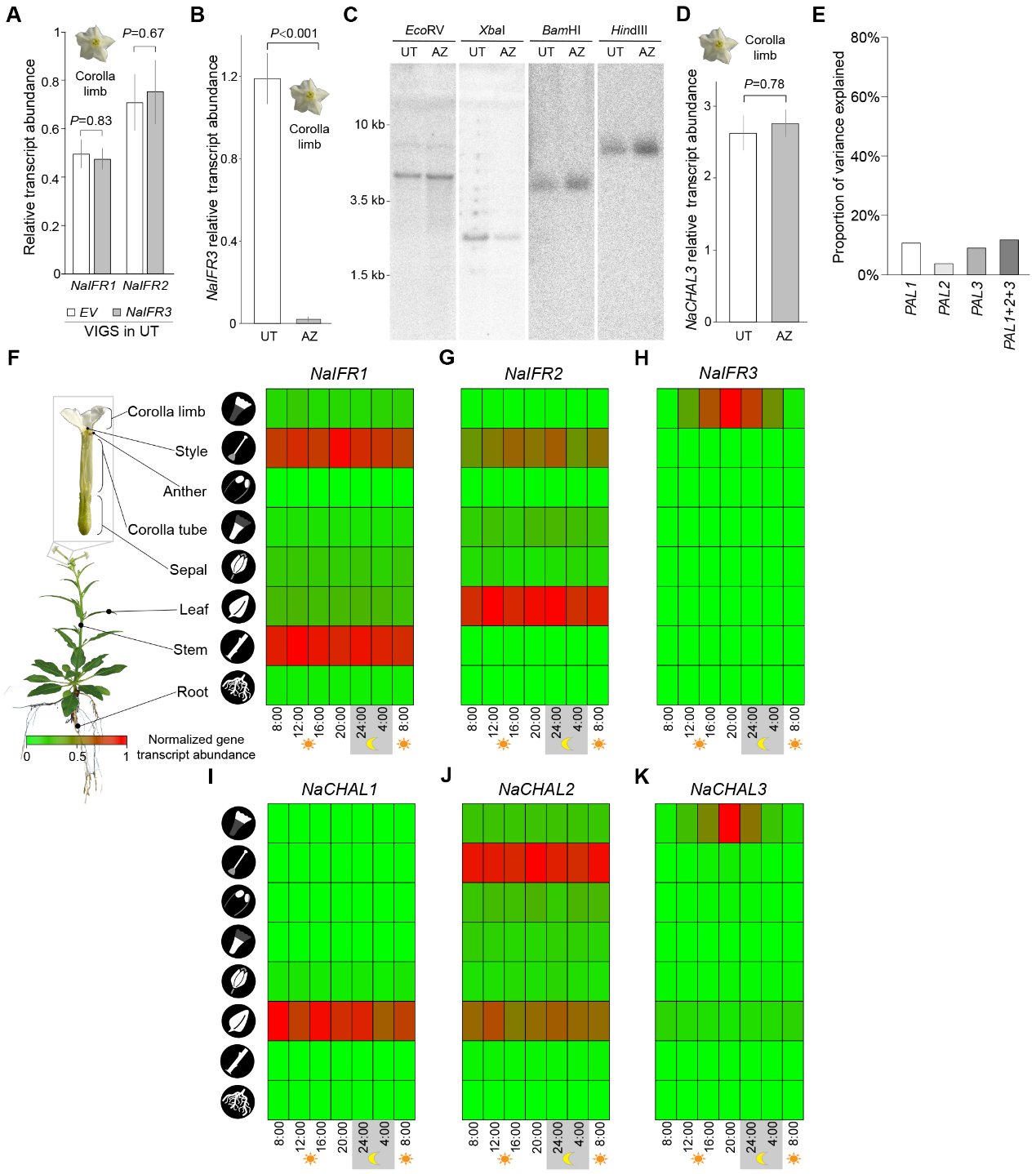

**Figure S2. Transcript abundance of *NaIFR* and *NaCHAL*.**

(A) Specificity of the *NaIFR3* silencing by VIGS in the corolla limb of the UT genotype. Corolla limb samples were harvested at 20:00. Transcript abundance (mean ± SE, n = 4) of *NaIFR1* and *NaIFR2* were analyzed by RT-qPCR and are shown relative to *NaEF*.

(B) Transcript abundance of *NaIFR3* (mean ± SE, n = 4) in the corolla limb is significantly different between UT and AZ genotypes. Corolla limb samples were harvested at 20:00. Transcript abundance was analyzed by RT-qPCR and is shown relative to *NaEF*.

(C) Genomic DNA (20 μg) from UT and AZ was digested with *Eco*RI or *Xba*I or *Bam*HI or *Hin*dIII, respectively. After separation by agarose gel electrophoresis and blotting, the DNA was hybridized with *NaIFR3* partial cDNA (453-692 bp, 240 bp). The blot was washed at high stringency (0.1× SSC and 0.5% SDS for 20 min twice).

(D) Transcript abundance of *NaCHAL3* in the corolla limb is not significantly different between UT and AZ genotypes. The corolla limb samples were harvested at 20:00. Transcript abundance (mean ± SE, n = 4) was analyzed by RT-qPCR and is shown relative to *NaEF*.

For (A), (B) and (D), *P* values were calculated using Student’s t tests.

(E) The y-axis denotes the proportion of BA variation, which could be explained by the changes of genes’ transcription.

(F) to (H), Heatmap representing the transcription kinetics of *NaIFR1* (F), *NaIFR2* (G) and *NaIFR3* (H) among eight tissues of UT genotype.

(I) to (K), Heatmap representing the transcription kinetics of *NaCHAL1* (I), *NaCHAL2* (J) and *NaCHAL3* (K) among eight tissues of UT genotype.

For (F) to (K), The transcript kinetics are shown relative to *NaEF* and normalized as X’ = X/X_max_ among different tissues.

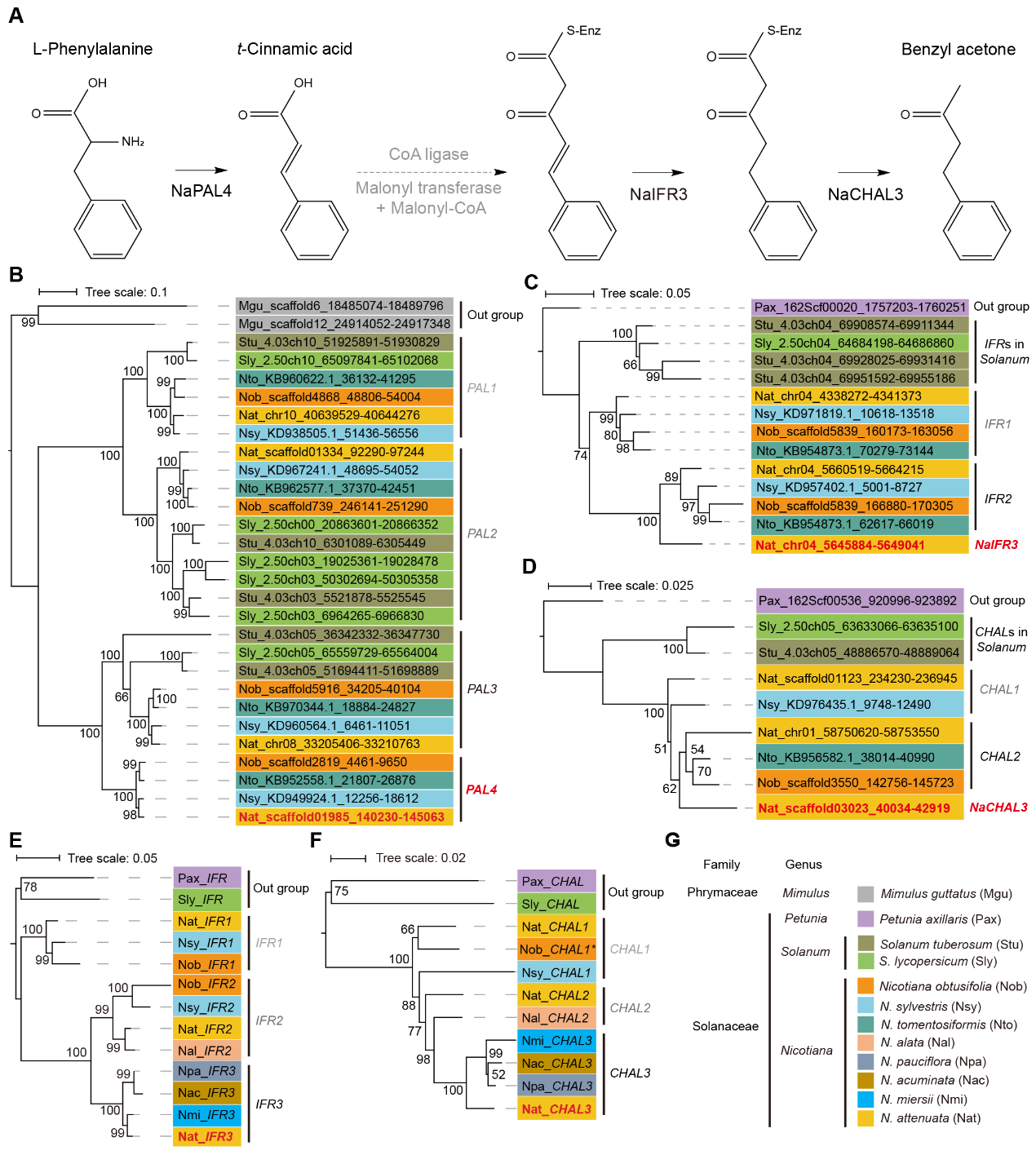

**Figure S3. Evolution of BA biosynthesis.**

(A) Predicted biosynthesis pathway of BA. Dashed lines with arrow heads indicate the putative steps and the putative corresponding enzymes are shown in grey.

(B) to (D), The phylogenetic trees of homologous genes of *PAL* (B), *IFR* (C) and *CHAL* (D) were built based on genomic data from the sequenced genomes. Numbers on branches indicate the bootstrap percentage values calculated from 100 replicates, and only values greater than 50% are shown. *PALs* in *Mimulus guttatus* were used as an outgroup for the phylogenetic analysis of *PAL* (B); *IFR* and *CHAL* in *Petunia axillaris* were used as an outgroup for the phylogenetic analysis of *IFR* (C) and *CHAL* (D), respectively. Sequence ID was shown as: species abbreviation _ genome version _ chromosome/scaffold ID _ positions of gene.

(E) and (F), Phylogenetic trees of the homologous genes of *IFR* (E) and *CHAL* (F) were built based on the cDNA sequences (except outgroups). Numbers on branches indicate the bootstrap percentage values calculated from 100 replicates, and only values greater than 50% are shown. *IFR* and *CHAL* in *Petunia axillaris* and *Solanum lycopersicum* were used as an outgroup for the phylogenetic analysis of *IFR* (E) and *CHAL* (F), respectively. In (F), *N. obtusifolia*_*CHAL1* (Marked with asterisk) was missing in its genomic database (likely due to incomplete genome assembly), but was identified by homologous cloning from *N. obtusifolia* corolla cDNA.

(G) Species used in the phylogenetic analysis in (B) to (F) with corresponding colors are shown.

| Primer ID | Sequence (5’ to 3’) | Purpose |
| --- | --- | --- |
| PAL1-F | caccATGGCCGGTGTTGCACAAAATGGTC | Intra- and interspecific homologous gene cloning (full-length CDS) |
| PAL1-R | ACAGATTGGTAGAGGAGCACCATTC |  |
| PAL2-F | caccATGGCATCAAATGGTCATGTTAATGGAG |  |
| PAL2-R | ACAGATAGGAAGAGGAGCACC |  |
| PAL3-F | caccATGGAGTATGCCAATGGAAATTGTAATG |  |
| PAL3-R | GCAAAGTGGAAGAGGAGCAC |  |
| PAL4-F | caccATGGAGTATGCAAATGGAGATTGTAATG |  |
| PAL4-R | GCAGAGTGGAAGAGGAGCACC |  |
| IFR3-F | caccATGGCTGAGAAAAGCAAAGTTCTGATC |  |
| IFR3-R | GACAAATTGAACAAGGCACTCCTCC |  |
| CHAL3-F | caccATGGTCACCGTCGAGGAGGTTCGAAG |  |
| CHAL3-R | AGTGGAGACACTATGAAGCACAACAGTCTC |  |
| PAL1-RT-F | TGTCAACGATTACTACAACAACGGG | Intraspecific RT-qPCR |
| PAL1-RT-R | TCAGATCCCTTGAAACCATAGTCCA |  |
| PAL2-RT-F | AACCAAAGCAAGATCGTTATGCTCT |  |
| PAL2-RT-R | TCTTAGTTGCAGAGCGAATGACTTC |  |
| PAL3-RT-F | AAGCCATTGACTTGAGACACTTGG |  |
| PAL3-RT-R | CACTTTCTTGGCCACTAAACTCACT |  |
| PAL4-RT-F | TGCAGTCACCAGGAGAAGTATTT |  |
| PAL4-RT-R | GCAATCAAGCAAAGGATCTATCAAT |  |
| IFR1-RT-F | CGTGGTAATATCAACATTGGGGATG |  |
| IFR1-RT-R | AACTTTCACAGCAAATGCAGACTTTGC |  |
| IFR2-RT-F | AGCAACTGGATGTATTGGAAAATATT |  |
| IFR2-RT-R | AGTAGTGTGCTTTCTCTAACCA |  |
| IFR3-RT-F | TATCGTTCAATGAGTTAGTGGCTA |  |
| IFR3-RT-R | AGGAGATGTATTGATGTCTTTGAGA |  |
| CHAL1-RT-F | ATGGTCACCGGCGAAAATC |  |
| CHAL1-RT-R | TAATAATCAGGATAGGTGCTTC |  |
| CHAL2-RT-F | CCAAAAGGCCATCAAAGAATGGGGC |  |
| CHAL2-RT-R | CCCGGGCATGTCCACTCCACTAG |  |
| CHAL3-RT-F | GCCACTCCTTCGAACTGTGTT |  |
| CHAL3-RT-R | AATCATTGATTTGTCACACATGCGC |  |
| EF-RT-F | CCACACTTCCCACATTGCTGT |  |
| EF-RT-R | CGCATGTCCCTCACAGCAAAAC |  |
| NaPAL1-VIGS-F | CTCTTGCCCCTTCAGAAAGCTCCACTG | VIGS |
| NaPAL1-VIGS-R | ATGGCCGGTGTTGCACAAAATGGTCACC |  |
| NaPAL2-VIGS-F | AAAGTTCAACTTTAACACCATTTGCACTTTT |  |
| NaPAL2-VIGS-R | ATGGCATCAAATGGTCATGTTAATGGAGG |  |
| NaPAL3-VIGS-F | TGAAGTGCTAGAATTCTTCTCCGCGTC |  |
| NaPAL3-VIGS-R | TCTCAAGGCTTCAGTGAAAAGTGCAGTG |  |
| NaPAL4-VIGS-F | TGAAGTGCTAGAAGTCTTCTCACCTTCACC | VIGS, Southern blot |
| NaPAL4-VIGS-R | TCTCAAGGCCTCAGTAAAAAATGTTGTGAG |  |
| NaIFR3-VIGS-F | TCCCACATAGCCACTAACTCATTGAACG |  |
| NaIFR3-VIGS-R | AAGCGACTTGTTAGATGGTTATTTCTTAGC | VIGS, Southern blot |

**Table S1. DNA primers used in this study.** Lowercase letters in the primer sequences indicate the 4 base pair sequences (CACC) necessary for directional cloning on the 5′ end of the forward primer.

| Gene or genome | Species | GenBank accession number |
| --- | --- | --- |
| *N. attenuata* genome of UT | *Nicotiana attenuata* | GenBank ID: MJEQ00000000 |
| *N. attenuata* genome of AZ | *Nicotiana attenuata* | GenBank ID: MCOF00000000 |
| Advanced Intercross RILs Genotyping by Sequencing *N. attenuata* | *Nicotiana attenuata* | GenBank ID: PRJNA378521 |
| *N. obtusifolia* genome | *Nicotiana obtusifolia* | GenBank ID: MCJB00000000 |
| *N. sylvestris* genome | *Nicotiana sylvestris* | GenBank ID: ASAF01000000 |
| *N. tomentosiformis* genome | *Nicotiana tomentosiformis* | GenBank ID: ASAG01000000 |
| *Mimulus guttatus* genome | *Mimulus guttatus* | GenBank ID: APLE01000000 |
| *Petunia axillaris* genome v1.6.2 | *Petunia axillaris* | <https://solgenomics.net/organism/Petunia_axillaris/genome> |
| *Solanum lycopersicum* genome v2.50 | *Solanum lycopersicum* | <https://solgenomics.net/organism/Solanum_lycopersicum/genome> |
| *Solanum tuberosum* genome v4.03 | *Solanum tuberosum* | <https://solgenomics.net/organism/Solanum_tuberosum/genome> |
| *NaPAL4* | *Nicotiana attenuata* | GenBank ID: XM019375988.1 |
| *NaIFR3* | *Nicotiana attenuata* | GenBank ID: XM019389443.1 |
| *NaCHAL3* | *Nicotiana attenuata* | GenBank ID: XM019379962.1 |
| *NaPAL1* | *Nicotiana attenuata* | GenBank ID: DQ768746.1 |
| *NaPAL2* | *Nicotiana attenuata* | GenBank ID: DQ768747.1 |
| *NaPAL3* | *Nicotiana attenuata* | GenBank ID: XM019396809.1 |
| *NaIFR1* | *Nicotiana attenuata* | GenBank ID: XM_019389418.1 |
| *NaIFR2* | *Nicotiana attenuata* | GenBank ID: XM_019389704.1 |
| *NaCHAL1* | *Nicotiana attenuata* | GenBank ID: XM_019392482.1 |
| *NaCHAL2* | *Nicotiana attenuata* | GenBank ID: XM_019370755.1 |
| *N. acuminataPAL4* | *Nicotiana acuminata* | GenBank ID: MK689220 |
| *N. acuminataIFR3* | *Nicotiana acuminata* | GenBank ID: MK689221 |
| *N. acuminataCHAL3* | *Nicotiana acuminata* | GenBank ID: MK689222 |
| *N. alataPAL4* | *Nicotiana alata* | GenBank ID: MK689223 |
| *N. miersiiPAL4* | *Nicotiana miersii* | GenBank ID: MK689229 |
| *N. miersiiIFR3* | *Nicotiana miersii* | GenBank ID: MK689230 |
| *N. miersiiCHAL3* | *Nicotiana miersii* | GenBank ID: MK689231 |
| *N. obtusifoliaCHAL1* | *Nicotiana obtusifolia* | GenBank ID: MK689234 |
| *N. paucifloraPAL4* | *Nicotiana pauciflora* | GenBank ID: MK689235 |
| *N. paucifloraIFR3* | *Nicotiana pauciflora* | GenBank ID: MK689236 |
| *N. paucifloraCHAL3* | *Nicotiana pauciflora* | GenBank ID: MK689237 |

**Table S2. Data deposition in this study.**

| Genotype | latitude | longitude |
| --- | --- | --- |
| UT | N37°19'36.26" | W113°57'53.05" |
| AZ | N35°12'56.07" | W111°27'41.29" |
| G2 | N37°04'33.53" | W113°49'58.74" |
| 43 | N37°19'35.48" | W113°57'38.28" |
| 84 | N37°19'36.48" | W113°57'33.28" |
| 85 | N37°21'07.103" | W114°05'50.298" |
| 97 | N37°21'35.24" | W113°56'38.68" |
| 133 | N37°06'12.5" | W113°49'36.6" |
| 138 | N37°8'19.58" | W114°1'35.10" |
| 149 | N35°12'56.07" | W111°27'41.29" |
| 176 | N37°16'38.65" | W113°53'35.18" |
| 179 | N37°21'1.04" | W113°57'5.17" |
| 194 | N37°20'22.52" | W114°2'40.86" |
| 278 | N37°21'02.580" | W114°05'53.661" |
| 304 | N37°20'22.52" | W114°2'40.86" |
| 305 | N37°45'19.61" | W118°35'41.82" |
| 308 | N37°13'5.50" | W113°48'24.25" |
| 331 | N37°13'15.83" | W113°48'20.86" |
| 341 | N37°9'45.30" | W114°0'58.52" |
| 351 | N37°17'09.1" | W114°07'31.5" |
| 370 | N37°9'1.30" | W113°47'43.36" |
| 382 | N37°14'27.05" | W113°49'36.71" |
| 384 | N37°14'27.05" | W113°49'36.71" |

**Table S3. GPS coordinates of the seed collection sites of the *N. attenuata* accessions.**
